# TRPML1 loss drives lysosomal calcium failure and astrocyte dysfunction across Alzheimer’s Disease progression

**DOI:** 10.64898/2026.07.23.740052

**Authors:** D. Shah, P. Desai, C. Chertavian, M.J. Thackray, M. Dumoulin, V.F.T. Mitchener, M. Strom, B. De Strooper, I.L Arancibia-Cárcamo

## Abstract

Astrocytes are among the earliest cells to exhibit dysfunction in Alzheimer’s disease (AD), developing profound calcium signalling deficits before amyloid plaques have formed, yet the underlying mechanisms remain unknown. Lysosomal dysfunction is a hallmark of AD, but whether it initiates this early functional impairment or arises as a consequence of established pathology remains unresolved. Here, we find that astrocytic cytosolic calcium activity is suppressed prior to amyloid plaque deposition and is accompanied by reduced lysosomal acidification *in vivo*. Using a lysosome-targeted calcium indicator selectively expressed in astrocytes, we directly visualise lysosomal calcium dynamics in vivo and reveal a profound early loss of lysosomal calcium release, identifying lysosomal failure as an initiating event in astrocyte dysfunction in AD. Reduced expression of the lysosomal calcium channel TRPML1 provides the mechanistic basis for this deficit. Astrocyte-specific restoration of TRPML1 expression rescues lysosomal homeostasis and cytosolic calcium signalling and prevents astrocyte reactivity and morphological hypertrophy. Strikingly, early TRPML1 restoration prevents both the initial calcium hypoactivity observed before plaque formation and the later hyperactivity that characterises post-plaque disease, demonstrating that lysosomal calcium homeostasis stabilises astrocyte function across the disease trajectory. TRPML1 restoration also reduces amyloid plaque burden, indicating that astrocytic lysosomal competence directly shapes disease pathology. These findings identify lysosomal calcium failure as an early organelle-level mechanism linking amyloid stress to astrocyte dysfunction in AD, and position TRPML1-mediated lysosomal calcium signalling as a tractable target for limiting disease progression.

## Introduction

Astrocytes are among the first cells to fail in Alzheimer’s disease (AD), emerging as active contributors to pathophysiology well before the onset of amyloid plaque deposition. Beyond their traditional supportive roles, astrocytes operate at the interface of neuronal activity, extracellular homeostasis, and circuit stability, coordinating these processes through tightly regulated calcium signalling^1^. Astrocytic calcium transients shape synaptic transmission, support neurovascular coupling, and tune metabolic support to neurons^2^. Disruptions of these dynamics therefore have the potential to affect neural circuit function broadly. Previous work demonstrated that astrocytic calcium activity is markedly reduced at very early stages of AD, before amyloid-β (Aβ) deposition, and that restoring this calcium deficit is sufficient to normalise neuronal activity, correct behavioural abnormalities, and recover network function^3^. These findings identify astrocytic dysfunction as one of the earliest detectable functional abnormalities in AD and suggest that early disturbances in astrocyte signalling have consequences extending beyond the glial compartment. However, the cellular machinery that destabilises astrocytic calcium homeostasis at these initial stages remains unknown.

More broadly, early AD pathology is increasingly understood as a failure of cellular homeostasis triggered by amyloid stress^4^, yet the subcellular mechanisms through which this stress is translated into functional impairment remain poorly defined. Lysosomes represent a compelling candidate in this context. Beyond their classical role in degradation, lysosomes function as central hubs that integrate metabolic state, proteostasis, and intracellular signalling, including calcium dynamics^5,6^. Through calcium dependent regulation of trafficking, membrane fusion, and adaptive responses to cellular stress, lysosomes are well positioned to couple amyloid stress to changes in cell function. Among the channels mediating this release, TRPML1 plays a central role in regulating lysosomal calcium signalling. By coupling calcium release to lysosomal trafficking, autophagy, and exocytosis, TRPML1 coordinates multiple aspects of lysosomal function and adaptive capacity^7^. Disruption of this pathway could therefore represent an early vulnerable point across brain cell types, with cell-type-specific downstream consequences. However, in AD, lysosomal dysfunction has largely been considered a downstream or plaque-associated phenomenon, and most studies in astrocytes have been restricted to Aβ-exposed cultures or plaque-proximal cells at later disease stages^8–10^, leaving the earliest *in vivo* response largely unexplored. Whether lysosomal calcium is disrupted as part of the earliest cellular response to amyloid stress in the intact brain and contributes to the initial destabilisation of astrocyte function *in vivo* remains unresolved.

Here we provide the first *in vivo* measurement of astrocytic lysosomal calcium dynamics in the brain. We show that lysosomal calcium release is impaired in astrocytes before plaque deposition in AD, and identify reduced TRPML1 expression as the mechanistic basis. Restoring TRPML1 selectively in astrocytes corrects the functional and structural trajectory of astrocyte dysfunction and reduces amyloid burden, linking astrocyte dysfunction to the emergence of pathological hallmarks of AD. These findings identify lysosomal calcium failure as an initiating subcellular event underlying astrocyte dysfunction in AD.

## Results

Astrocytic calcium signalling is markedly reduced before amyloid-β deposition becomes detectable, a functional deficit with broad consequences for neural circuit homeostasis ^3^. To identify the cell-intrinsic mechanisms responsible, we examined heterozygous *App^NL-G-F^* mice (*App^WT/NL-G-F^)*, which develop amyloid plaques at 6–7 months of age^11^, enabling analysis in adult mice at clearly defined pre-plaque (3 months) and post-plaque stages (8 months) (Fig. S1). Consistent with our previous findings, *in vivo* two-photon imaging revealed a reduction in cytosolic calcium activity in 3-month-old *App^WT/NL-G-F^* mice relative to wild-type controls. Both transient amplitude (ΔF/F₀: 0.915 vs. 0.217, *p=*0.0311) and frequency (peaks/min: 0.144 vs. 0.057, *p=*0.004) were strongly suppressed (Fig. 1A), confirming an early and robust loss of astrocytic calcium signalling before detectable plaque deposition. Given their role as dynamic intracellular calcium stores and their sensitivity to metabolic and proteostatic stress^12,13^, we next examined whether astrocytic lysosomes show early functional abnormalities that could account for this signalling deficit. Using FIREpHLy, a ratiometric pH sensitive probe targeted to the lysosomal lumen and selectively expressed in astrocytes under the *GfaABC1D* promoter, we found that astrocytic lysosomes in 3-month-old *App^WT/NL-G-F^* mice are significantly less acidic than in wild-type controls (mTFP/mCherry ratio WT vs. *App^WT/NL-G-F^*: 0.58 vs. 0.72; *p* < 0.001; Fig. 1E). Moreover, these results were accompanied by structural alterations in astrocyte lysosomes. Immunostaining showed significant increases in the lysosomal markers (WT vs. *App^WT/NL-G-F^*) LAMP1 (% area 0.84 vs. 1.64, *p=*0.0001), Cathepsin D (% area 1.65 vs. 2.64, *p=*0.0028), and LAMP2 (% area 1.78 vs. 2.38, *p=*0.02) (Fig. 1C, Fig. S2) in AD astrocytes, consistent with abnormal lysosomal accumulation and enlargement. Together, these data show that astrocytic lysosomes display both structural and functional abnormalities before plaque deposition.

**Figure 1:**
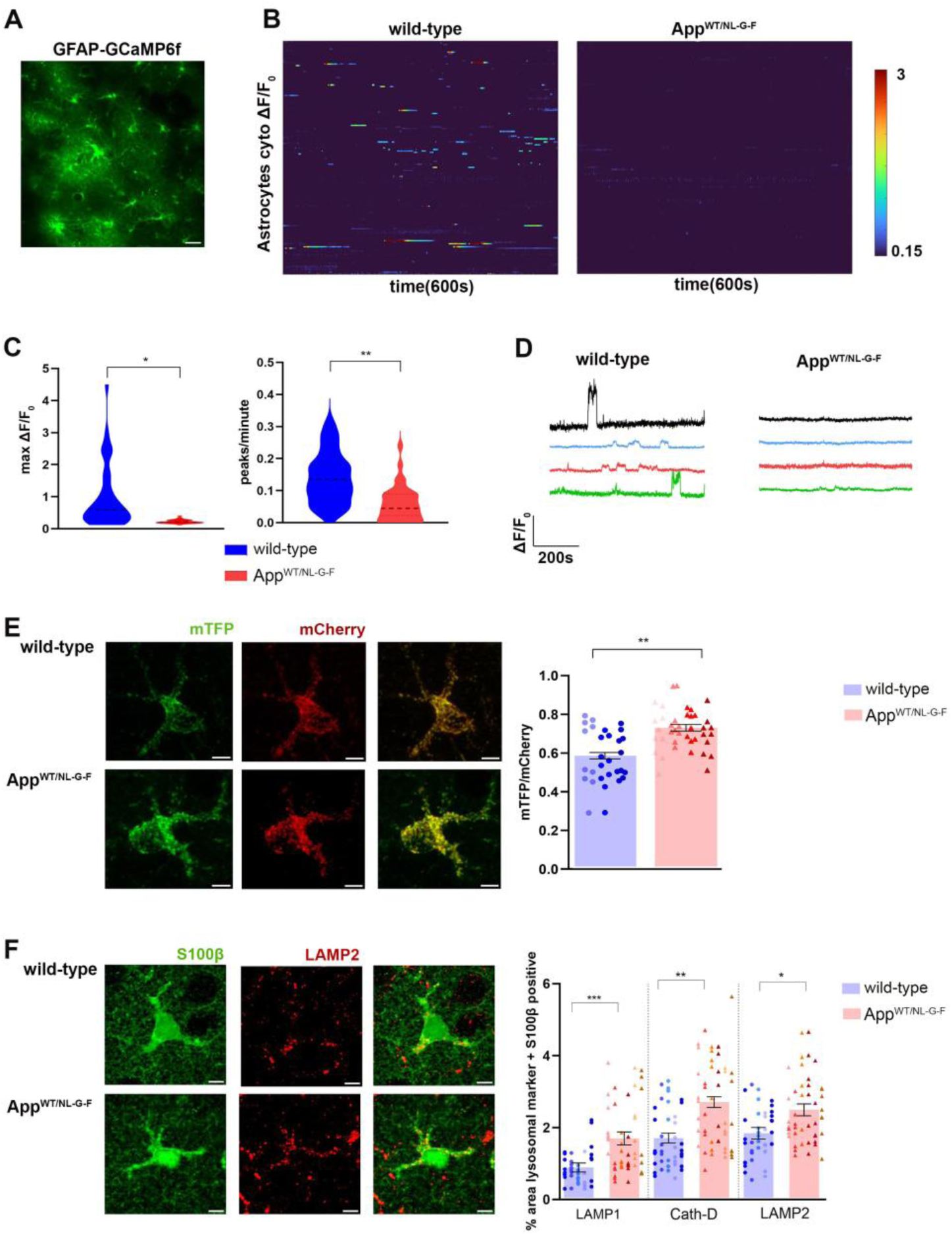
Early cytosolic calcium dynamics and abnormalities of lysosomal function in AD astrocytes. A) *In vivo* two-photon calcium imaging following AAV-GFAP-GCaMP6f expression. B) Heatmaps display cytosolic calcium transients (ΔF/F₀) across astrocytes over a 600 s period. C) Quantification of ÄF/F₀ amplitude and frequency (peaks/min) is shown for wild-type and App^WT/NL-G-F^ mice (*N* = 4/group, 27–34 astrocytes per mouse; mean ± SEM). Amplitude: 0.915 ± 0.14 (WT) vs. 0.217 ± 0.01 (App^WT/NL-G-F^), *p* = 0.0311; frequency: 0.144 ± 0.012 (WT) vs. 0.057 ± 0.009 (App^WT/NL-G-F^), *p =* 0.004. Statistical test: ART ANOVA. **D)** Representative calcium time traces are shown for each experimental group. **E)** Representative confocal images showing expression of the AAV-GFAP-FIRE**-**pHLy probe in astrocytes of WT (*N* = 3 mice, 10 cells per mouse) and App^WT/NL-G-F^ mice (*N* = 4 mice, 10 cells per mouse). Quantification of the mTFP/mCherry ratio (mean ± SEM): 0.58 ± 0.02 (WT) vs. 0.72 ± 0.01 (App^WT/NL-G-F^)*, p = 0.0042.* Statistical test: Nested two-sample *t*-test. **F)** Immunohistochemistry images showing LAMP1, Cathepsin D and LAMP2 expression in astrocytes (S100β) from wild-type (WT; *N* = 5 mice, 8 cells per mouse) and App^WT/NL-G-F^ mice (*N* = 6 mice, 8 cells per mouse). Quantification of LAMP1, Cathepsin D and LAMP2 is shown as % area of pixels double-positive for each lysosomal marker and S100β (mean ± SEM). WT vs. App^WT/NL-G-F^ values: LAMP1, 0.84 ± 0.12 vs. 1.64 ± 0.18 (*p* = 0.0001); Cathepsin D, 1.65 ± 0.14 vs. 2.64 ± 0.15 (*p* = 0.0028); LAMP2, 1.78 ± 0.16 vs. 2.38 ± 0.18 (*p* = 0.02). Statistical test: One-way ANOVA with Holm-Sidak correction.

Importantly, these findings establish lysosomal dysfunction as an early event in AD astrocytes, but it is not known whether these impairments extend to functional changes in lysosomal calcium release and to what extent they contribute to the cytosolic calcium disruption. Lysosomal calcium dynamics have not previously been measured *in vivo* in the brain. To address this gap, we expressed a LAMP1-targeted GCaMP6f probe selectively in astrocytes to directly measure lysosomal calcium release in the living brain. In wild-type mice, this approach revealed robust spontaneous lysosomal calcium transients in astrocytes, confirming that lysosomal calcium release is a dynamic and measurable feature of astrocyte physiology *in vivo*. Importantly, our data show that lysosomal calcium peaks were almost abolished in astrocytes of 3-month-old *App^WT/NL-G-F^* mice. Both the amplitude (WT vs. *App^WT/NL-G-F^* ΔF/F₀: 1.85 vs. 0.57, *p=*0.0023) and frequency (WT vs. *App^WT/NL-G-F^* peaks/min: 0.27 vs. 0.068, *p=*0.0036) of lysosomal calcium transients were strongly suppressed compared to wild-type controls (Fig. 2A) indicating a substantial loss of lysosomal Ca2+ release.

**Figure 2:**
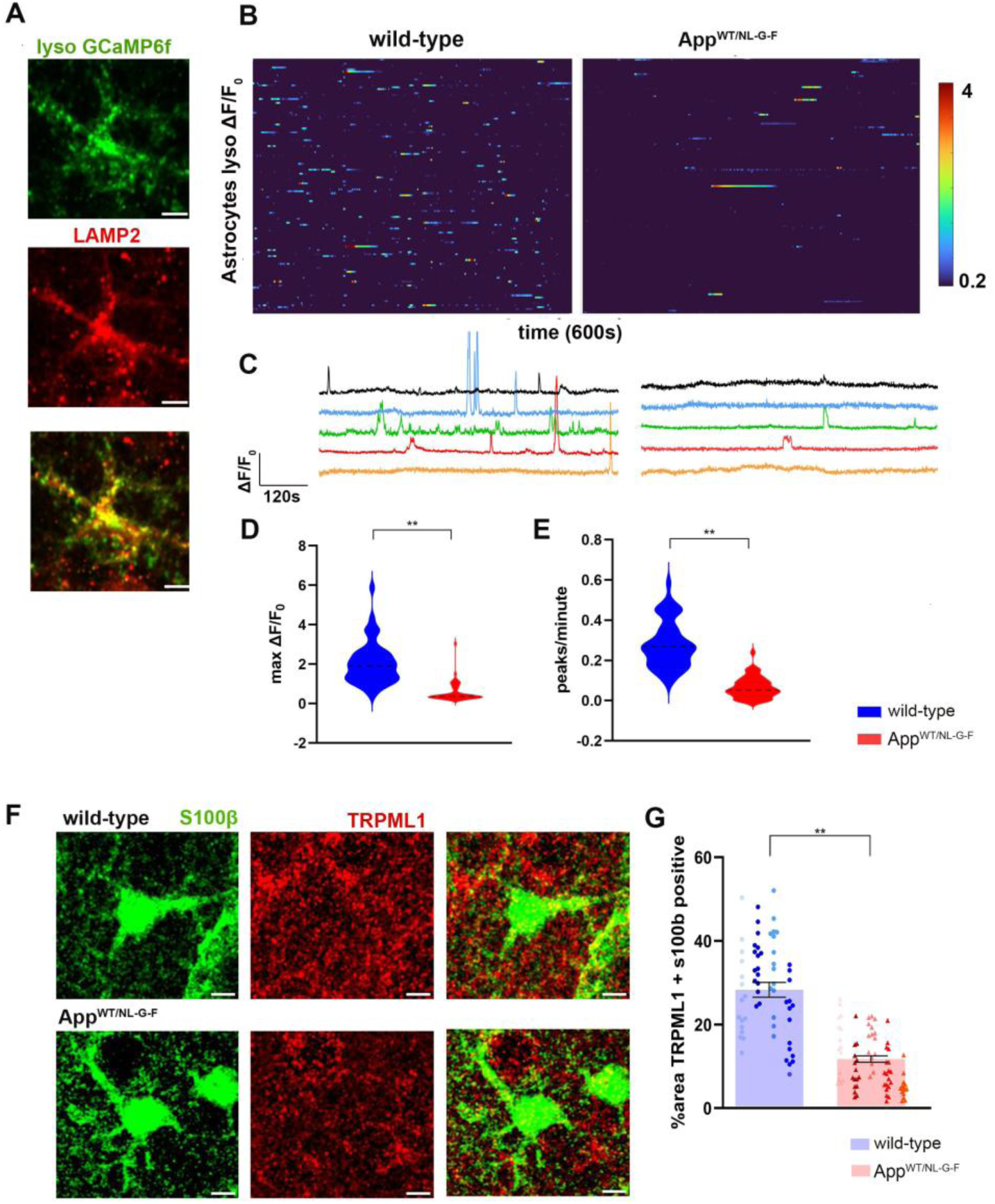
Reduced lysosomal calcium release in AD astrocytes. **A)** Immunohistochemistry images showing expression of the AAV-GFAP-LAMP1-GCaMP6f probe, with co-localisation of LAMP2 confirming lysosomal targeting. **B-C)** *In vivo* two-photon calcium imaging following AAV-GFAP-LAMP1-GCaMP6f expression. Heatmaps depict lysosomal calcium transients (ΔF/F₀) over 600 s for individual astrocytes; five representative traces are shown below for each group. **D-E)** Quantification of ΔF/F₀ amplitude and event frequency (mean ± SEM) is shown for WT and App^WT/NL-G-F^ mice (*N* = 4 mice/group, 25–40 astrocytes per mouse). Amplitude: 1.85 ± 0.15 (WT) vs. 0.57 ± 0.07 (App^WT/NL-G-F^), *p* = 0.0023. Frequency: 0.27 ± 0.018 (WT) vs. 0.068 ± 0.009 (App^WT/NL-G-F^), *p* = 0.0036. Statistical test: ART ANOVA. **F-G)** Immunohistochemistry images showing TRPML1 expression in astrocytes (S100β) from wild-type (WT; *N* = 4 mice, 15–16 cells per mouse) and App^WT/NL-G-F^ mice (*N* = 5 mice, 15–19 cells per mouse). Quantification of % area double-positive for TRPML1 and S100β (mean ± SEM): WT, 29.04 ± 1.36; App^WT/NL-G-F^, 10.61 ± 0.72; *p* = 0.003. Statistical test: ART ANOVA.

We next sought the molecular basis for this deficit. TRPML1 has been identified as the principal Ca2+ permeable channel on the lysosomal membrane and a key regulator of lysosomal function and integrity^7,12^. Consistent with this role, selective overexpression of TRPML1 in astrocytes was sufficient to enhance lysosomal calcium release in wild-type mice *in vivo* (Fig. S3), directly demonstrating that TRPML1 controls lysosomal calcium dynamics in astrocytes in the intact brain. Strikingly, immunohistochemistry revealed that TRPML1 expression was substantially reduced in astrocytes from *App^WT/NL-G-F^* mice compared to wild-type (% area WT vs. *App^WT/NL-G-F^* 29.04 vs. 10.61, *p=*0.003) (Fig. 2B). Together, these data identify reduced TRPML1 expression as a mechanistic basis for the lysosomal Ca2+ deficit observed *in vivo*.

Having identified loss of TRPML1-mediated lysosomal calcium release as a driver of early astrocytic signalling deficits, we next asked whether restoring TRPML1 expression could rescue cytosolic calcium dynamics *in vivo*, and whether this intervention would also correct the underlying lysosomal dysfunction. We overexpressed TRPML1 selectively in astrocytes of *App^WT/NL-G-F^* mice and performed 2-photon calcium imaging. Consistent with our earlier findings (Fig. 1), astrocytes in *App^WT/NL-G-F^* mice expressing an mCherry control construct exhibited the previously shown AD-associated suppression of cytosolic calcium activity, with transient amplitude suppressed by ∼75% and frequency reduced by ∼43% relative to wild-type mCherry controls (ΔF/F₀ *p* = 0.0093 and peaks/min *p* = 0.010; Fig. 3).

**Figure 3:**
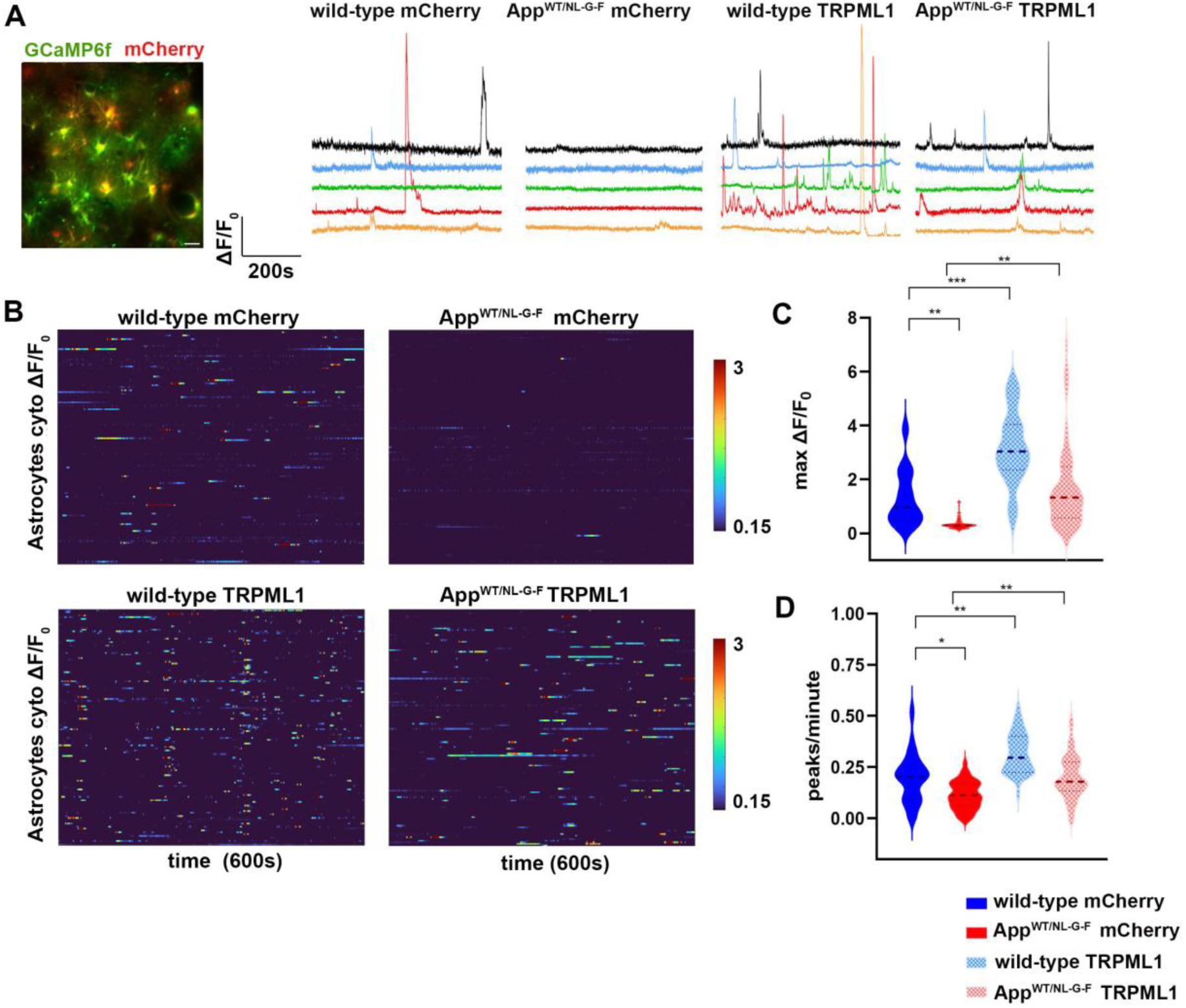
Overexpression of TRPML1 rescues cytosolic calcium signals in astrocytes. **A)** *In vivo* two-photon calcium imaging following co-expression of AAV-GFAP-GCaMP6f with either AAV-GFAP-mCherry (control) or AAV-GFAP-TRPML1. Five representative time courses are shown for each experimental group. **B)** Heatmaps depict cytosolic calcium transients (ΔF/F₀) over 600 s for individual astrocytes. **C-D)** Quantification of ΔF/F₀ amplitude and frequency (mean ± SEM) is shown for wild-type (WT) mCherry controls (*N* = 4 mice, 32–37 astrocytes per mouse), App^WT/NL-G-F^ mCherry controls (*N* = 4 mice, 30–39 astrocytes per mouse), WT TRPML1 (*N* = 4, 31–35 astrocytes per mouse), and App^WT/NL-G-F^ TRPML1 (*N* = 4 mice, 33–39 astrocytes per mouse). **Amplitude:** WT mCherry, 1.37 ± 0.18; App^WT/NL-G-F^ mCherry, 0.34 ± 0.22; WT TRPML1, 3.17 ± 0.03; App^WT/NL-G-F^ TRPML1, 1.83 ± 0.21. Significant differences: WT mCherry vs. App^WT/NL-G-F^ mCherry, *p =* 0.0093; WT mCherry vs. WT TRPML1, *p =* 0.0006; App^WT/NL-G-F^ mCherry vs. App^WT/NL-G-F^ TRPML1, *p =* 0.0012. **Frequency:** WT mCherry, 0.198 ± 0.015; App^WT/NL-G-F^ mCherry, 0.113 ± 0.009; WT TRPML1, 0.314 ± 0.018; App^WT/NL-G-F^ TRPML1, 0.205 ± 0.014. Significant differences: WT mCherry vs. App^WT/NL-G-F^ mCherry, *p* = 0.010; WT mCherry vs. WT TRPML1, *p* = 0.009; App^WT/NL-G-F^ mCherry vs. TRPML1, *p* = 0.0034. Statistical test: ART ANOVA with Holm-Sidak correction.

Interestingly, TRPML1 overexpression substantially increased cytosolic calcium signalling in both genotypes. In wild-type astrocytes, it boosted amplitude by more than twofold (WT mCherry vs. TRPML1 ΔF/F₀: 1.37 vs. 3.17, *p=*0.0006) and increased frequency by ∼60% (WT mCherry vs. TRPML1 peaks/min: 0.198 vs. 0.314, *p=*0.009). Importantly, in *App^WT/NL-G-F^* astrocytes, TRPML1 overexpression restored cytosolic calcium activity to levels comparable to those measured in wild-type mCherry controls, significantly increasing both amplitude (*App^WT/NL-G-F^* mCherry vs. TRPML1 ΔF/F₀: 0.34 vs. 1.83, *p=*0.0012) and event frequency (*App^WT/NL-G-F^* mCherry vs. TRPML1 peaks/min: 0.113 vs. 0.205, *p=*0.0034). Together, these findings indicate that TRPML1-dependent lysosomal calcium release is a key determinant of cytosolic calcium homeostasis and that increasing TRPML1 expression is sufficient to rescue the signalling deficit in AD astrocytes (Fig. 3).

Notably, the effect of TRPML1 overexpression extended beyond cytosolic calcium signalling; TRPML1 expression normalised lysosomal acidity, as measured with the ratiometric FIRE-pHLy probe (Fig. S4), and reduced the elevated levels of the lysosomal marker LAMP2 in astrocytes of *App^WT/NL-G-F^* mice (Fig. S5). These findings indicate that restoring lysosomal calcium release not only rescues astrocytic calcium signalling but also re-establishes lysosomal homeostasis in AD astrocytes. Together, these data suggest that TRPML1 acts upstream of both lysosomal and cytosolic calcium signalling defects in AD astrocytes, placing it at the root of astrocytic dysfunction in AD.

Having demonstrated that TRPML1-mediated lysosomal calcium failure drives early astrocytic dysfunction, we next asked whether this initiating defect determines the subsequent trajectory of astrocyte signalling across disease progression. In mouse models of amyloid pathology, astrocytic calcium activity follows a well-described biphasic pattern: an initial period of marked hypoactivity in the pre-plaque stage^3,14^, followed by pronounced hyperactivity in the later stages of pathology^15^. Consistent with this, we show that astrocytes in 8-month-old *App^WT/NL-G-F^* mice displayed pronounced calcium hyperactivity, with transient amplitude increased ∼3.3-fold (WT vs. *App^WT/NL-G-F^* mCherry ΔF/F₀: 0.67 vs. 2.22, p<0.0001) and frequency increased ∼5-fold (WT vs. *App^WT/NL-G-F^*mCherry peaks/min: 0.131 vs. 0.683, p<0.0001) compared to wild-type controls (Fig. 4A-C).

**Figure 4:**
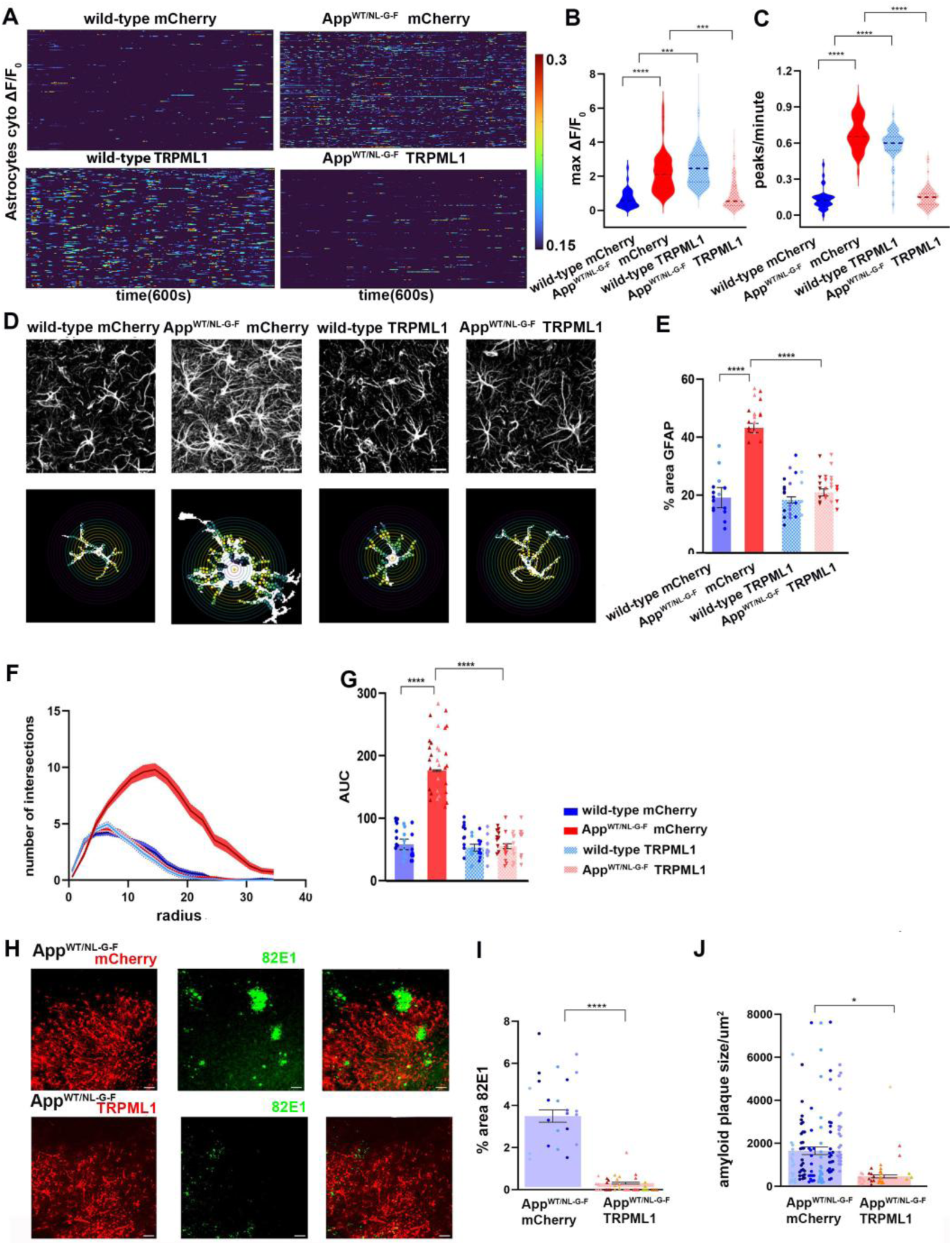
Early TRPML1 overexpression prevents late-stage astrocytic calcium hyperactivity and reduces plaque burden. **A)** *In vivo* two-photon calcium imaging following co-expression of AAV-GFAP-GCaMP6f with either AAV-GFAP-mCherry (control) or AAV-GFAP-TRPML1. Heatmaps depict cytosolic calcium transients (ΔF/F₀) over a 600 s imaging period for individual astrocytes. **B-C)** Quantification of ΔF/F₀ amplitude and frequency (mean ± SEM) is shown for wild-type (WT) mCherry controls (*N* = 4 mice, 32–36 astrocytes per mouse), App^WT/NL-G-F^ mCherry controls (*N* = 4 mice, 41–48 astrocytes per mouse), WT TRPML1 (*N* = 4 mice, 25–42 astrocytes per mouse), and App^WT/NL-G-F^ TRPML1 (*N* = 4 mice, 30–46 astrocytes per mouse). **Amplitude:** WT mCherry, 0.67 ± 0.07; App^WT/NL-G-F^ mCherry, 2.22 ± 0.14; WT TRPML1, 2.61 ± 0.18; App^WT/NL-G-F^ TRPML1, 0.87 ± 0.11. Significant differences: WT mCherry vs. App^WT/NL-G-F^ mCherry, *p* < 0.0001; WT mCherry vs. WT TRPML1, *p* = 0.0004; App^WT/NL-G-F^ mCherry vs. TRPML1, *p* = 0.0001. **Frequency:** WT mCherry, 0.131 ± 0.012; App^WT/NL-G-F^ mCherry, 0.683 ± 0.020; WT TRPML1, 0.609 ± 0.017; App^WT/NL-G-F^ TRPML1, 0.145 ± 0.012. Significant differences: WT mCherry vs. App^WT/NL-G-F^ mCherry, *p* < 0.0001; WT mCherry vs. WT TRPML1, *p* < 0.0001; App^WT/NL-G-F^ mCherry vs. TRPML1, *p* < 0.0001. Statistical test: ART ANOVA with Holm-Sidak correction. **D)** Upper panel shows representative GFAP immunostaining from wild-type (WT) and App^WT/NL-G-F^ mice injected with either AAV-GFAP-mCherry (control) or AAV-GFAP-TRPML1. Lower panel shows representative astrocytes with Sholl analysis circles illustrating morphological complexity. **E)** Quantification of astrocyte reactivity as % GFAP-positive area (mean ± SEM). Each color represents different mice (N = 3-4 mice per experimental group and 4-8 images/mouse). WT mCherry, 18.26 ± 1.66; App^WT/NL-G-F^ mCherry, 43.34 ± 1.25; WT TRPML1, 17.85 ± 1.27; App^WT/NL-G-F^ TRPML1, 21.27 ± 0.91. Significant differences: WT mCherry vs. App^WT/NL-G-F^ mCherry, *p* < 0.0001; App^WT/NL-G-F^ mCherry vs. TRPML1, *p* < 0.0001. Statistical test: nested one-way ANOVA with Holm-Sidak correction. **F)** Sholl profiles showing the number of intersections ± SEM. **G)** Area under the curve (AUC) from Sholl analysis (mean ± SEM). Each color represents different mice (N=3-4 mice per experimental group, 10 astrocytes per mouse). WT mCherry, 58.10 ± 3.95; App^WT/NL-G-F^ mCherry, 176.2 ± 8.02; WT TRPML1, 52.95 ± 2.95; App^WT/NL-G-F^ TRPML1, 55.25 ± 3.18. Significant differences: WT mCherry vs. App^WT/NL-G-F^ mCherry, *p* < 0.0001; App^WT/NL-G-F^ mCherry vs. TRPML1, *p* < 0.0001. Statistical test: nested one-way ANOVA with Holm-Sidak correction. **H-J)** Immunohistochemistry images showing 82E1 staining for amyloid-β in regions of AAV expression. Quantification of amyloid burden is shown as % area positive for 82E1 staining (mean ± SEM) for App^WT/NL-G-F^ mCherry controls (*N* = 6 mice, 3–5 images per mouse) and App^WT/NL-G-F^ TRPML1 (*N* = 7 mice, 4–7 images per mouse). **Plaque area (%):** App^WT/NL-G-F^ mCherry, 3.87 ± 0.35 vs. TRPML1, 0.20 ± 0.05; *p* = 0.00036. **Plaque size (µm²):** App^WT/NL-G-F^ mCherry, 1728 ± 144 vs. TRPML1, 596 ± 127; *p* = 0.036. Statistical test: ART ANOVA.

Remarkably, early restoration of TRPML1 expression attenuated this pathological shift. *App^WT/NL-G-F^* mice in which TRPML1 was expressed from 2 months of age did not develop the late-stage astrocytic hyperactivity characteristic of AD astrocytes in post-plaque environments. At 8 months, calcium signals in *App^WT/NL-G-F^* mice expressing TRPML1 remained close to wild-type levels (WT mCherry vs. *App^WT/NL-G-F^* TRPML1 ΔF/F₀: 0.67 vs. 0.87, *p=*0.28; peaks/min: 0.131 vs. 0.145, *p=*0.38), in contrast to the exaggerated activity of *App^WT/NL-G-F^*astrocytes expressing mCherry (*App^WT/NL-G-F^*mCherry vs. TRPML1 ΔF/F₀: 2.22 vs. 0.87 ± 0.11, *p*=0.0001; peaks/min: 0.683 vs. 0.145, *p*<0.0001) (Fig. 4A-C). In wild-type mice, TRPML1 continued to boost calcium activity above baseline (*p* = 0.0004 and *p* < 0.0001), consistent with its role in enhancing lysosomal and cytosolic Ca2+ signalling. Together, these findings indicate that early TRPML1 expression shapes disease trajectory by maintaining stable calcium homeostasis throughout progression.

Astrocyte reactivity, characterised by morphological hypertrophy and upregulation of GFAP, is a hallmark of AD pathology that emerges alongside plaque deposition. We therefore asked whether early TRPML1 restoration, by correcting the upstream lysosomal defect, could prevent this defining structural transformation. In *App^WT/NL-G-F^* mCherry mice, GFAP immunoreactivity was markedly elevated, and astrocyte morphology showed hypertrophy, with Sholl profiles shifted outward and increased AUC (Fig. 4D-G). Interestingly, early TRPML1 expression prevented these changes: GFAP coverage returned toward wild-type levels and Sholl/AUC metrics were normalised in *App^WT/NL-G-F^* TRPML1 mice (Fig. 4D-G). Early TRPML1 restoration thus prevented not only calcium hyperactivity but also the morphological hallmarks of astrocyte reactivity, suggesting that lysosomal calcium homeostasis regulates the reactive transformation of astrocytes in AD.

Together, these findings show that correcting lysosomal calcium release at an early stage stabilises astrocytic function and structure throughout disease progression. Therefore, we next asked whether this intervention also impacts core AD pathology. Quantification of 82E1 immunostaining in *App^WT/NL-G-F^* mice revealed a striking decrease in total plaque area (% area *App^WT/NL-G-F^* mCherry vs. TRPML1: 3.87 vs. 0.20, *p* = 0.00036). Interestingly, individual plaques were approximately 66% smaller in size (plaque size µm² in *App^WT/NL-G-F^* mCherry vs. TRPML1: 1728 vs. 596, *p* = 0.036; Fig. 4H-J). Importantly, the reduction in amyloid burden was confined to regions expressing TRPML1, indicating a local effect of astrocyte-specific modulation. Thus, early restoration of lysosomal calcium release not only corrects astrocytic hypoactivity (Fig. 3) but also prevents the emergence of late-stage hyperactivity and reactive transformation, and attenuates amyloid accumulation (Fig. 4), demonstrating that a single astrocyte-specific, organelle-level intervention can reshape the trajectory of AD pathology.

## Discussion

Here we identify lysosomal calcium failure as an initiating organelle-level mechanism driving astrocytic dysfunction in Alzheimer’s disease. This defect arises before amyloid plaque deposition and sits upstream of the functional and structural changes that characterise astrocyte impairment. Our findings advance understanding beyond the observation that astrocytic calcium is disrupted in early AD, providing a cell-intrinsic mechanism and a molecular entry point, TRPML1, through which amyloid stress is translated into astrocyte dysfunction. Although identified in astrocytes, this mechanism may reflect a broader vulnerability of cellular homeostasis in AD, whereby disruption of lysosomal signalling represents an early response to amyloid exposure.

We find that astrocytic lysosomes are structurally and functionally compromised in AD, with reduced acidification, increased marker accumulation, and impaired calcium release. These alterations are consistent with reduced lysosomal competence, which would be expected to affect proteostasis, metabolic adaptation, and intracellular signalling^16^. Importantly, these defects precede detectable plaque pathology, indicating that lysosomal dysfunction is an early cellular response to amyloid stress rather than a downstream feature of established pathology.

Lysosomal calcium signalling regulates membrane trafficking, organelle crosstalk, and ER calcium homeostasis^5,6,13,16^, making it a key integrator of intracellular signalling. We find that lysosomal calcium release is markedly reduced in astrocytes at early disease stages. This provides a mechanism by which lysosomal stress can propagate into broader cellular dysfunction. In astrocytes, this manifests as suppression of cytosolic calcium activity. Whether similar perturbations in lysosomal calcium handling contribute to dysfunction in other cell types, including neurons and microglia, remains to be determined.

We find that reduced TRPML1 expression provides the mechanistic basis for lysosomal calcium failure in AD astrocytes. TRPML1 is the principal lysosomal calcium release channel and plays an important role in maintaining lysosomal integrity, trafficking, and organelle crosstalk^7,16–18^. Critically, restoring TRPML1 is sufficient to improve both lysosomal function and cytosolic calcium signalling, demonstrating that TRPML1 loss is not merely associated with lysosomal dysfunction but drives it. Notably, this rescue extends beyond calcium release itself: TRPML1 restoration corrects impaired lysosomal acidification and accumulation, placing TRPML1 upstream of general lysosomal competence. Together, these findings identify TRPML1 as a regulator of lysosomal homeostasis and cellular signalling under stress conditions.

The consequences of this early lysosomal defect extend beyond the initial signalling phenotype. Astrocytes in AD transition from early hypoactivity to later hyperactivity and reactive remodelling^15^, a biphasic pattern that has been difficult to explain mechanistically. Our data are consistent with a model in which this progression arises from an upstream defect in lysosomal calcium signalling. Early loss of lysosomal calcium signalling suppresses astrocytic activity, whereas subsequent compensatory or pathological responses may drive hyperactivity and reactivity. The ability of TRPML1 restoration to attenuate both phases indicates that lysosomal calcium homeostasis acts as a stabilising mechanism that maintains astrocytes within a physiological range.

We find that astrocyte-specific restoration of TRPML1 also reduces amyloid plaque burden, demonstrating that early correction of astrocyte dysfunction is sufficient to limit disease progression. This finding argues against a purely reactive role for astrocytes in AD and instead supports a model in which early astrocyte dysfunction contributes causally to amyloid pathology. The precise mechanisms linking astrocytic calcium signalling to amyloid processing remain to be defined, but restored lysosomal competence may enhance astrocytic degradation and clearance of amyloid species through cell-intrinsic mechanisms. In parallel, normalisation of astrocytic calcium signalling reduces neuronal hyperactivity^3^ which may in turn limit amyloid production and amyloid accumulation in the local microenvironment. Beyond these effects, restoration of astrocyte homeostasis may influence glial crosstalk, including microglial surveillance and motility, processes that are critical for amyloid clearance. Together, these observations indicate that astrocyte homeostasis is a key determinant of the local amyloid microenvironment.

More broadly, our findings raise the possibility that lysosomal calcium dysfunction represents a shared early response to amyloid stress across multiple brain cell types^19–21^. Astrocytes, with their sensitivity to metabolic and proteostatic stress, may provide a particularly early readout of this process, but the same mechanism may drive distinct downstream phenotypes in neurons and microglia.

Together, this work identifies lysosomal calcium signalling as an early organelle-level mechanism linking amyloid stress to astrocyte dysfunction in vivo. By establishing TRPML1-dependent lysosomal calcium release as a key regulator of astrocyte homeostasis, our findings define a mechanistic pathway through which early cellular stress is translated into functional impairment. These results position lysosomal calcium signalling not as a downstream consequence of pathology, but as a point of control in the early stages of disease, with implications for therapeutic intervention prior to irreversible damage.

## Methods

### Mice

Male and female mice used in this study were bred and housed at the Francis Crick Institute under specific pathogen-free conditions. Heterozygous *App^NL-G-F^* knock-in mice, which carry the KM670/671N Swedish, E693G Arctic, and I716F Iberian mutations in the humanised *App* gene, were generated on a C57BL/6J background as described by Saito et al. (2014)^11^. Wild-type littermates were used as controls throughout the study. Heterozygous *APP^NL-G-F^* mice show the first cortical amyloid plaques around 6-7 months of age^11^. Mice were housed in individually ventilated cages with ad libitum access to food and water. All experimental procedures complied with the UK Animals (Scientific Procedures) Act and were approved by the relevant local ethical review board.

### Viral injections and Craniotomies

All AAVs used in this study were packaged in-house by the Vector Core Facility at the Francis Crick Institute at a titre of 1×10¹² GC/ml and serotyped as AAV2/8. For all constructs, the GFAP promoter refers to the truncated *GfaABC1D* variant, optimised for AAV-mediated expression^22^. The plasmids for pAAV-GFAP-GCaMP6f, pAAV-GFAP-LAMP1-GCaMP6f, pAAV-GFAP-TRPML1-mCherry, pAAV-GFAP-mtagBFP2 and pAAV-GFAP-TRPML1-mtagBFP2 were generated by the Francis Crick Vector Core. Two additional plasmids, pAAV-GFAP-mCherry (Addgene #58909) and the FIRE-pHLy probe (Addgene #170775), were obtained from Addgene. AAVs were produced by triple transfection of HEK-293T cells essentially as described in Challis et al.^23^ with the following exceptions: 3 x 10 cm dishes were used and after the PEG precipitation samples were resuspended in 4 ml GB +50U/ml Denarase incubated for 45 min at 37°C, then 2.5 ml Chloroform was added and the sample was vortexed for 2 min followed by centrifugation at 3000 x g for 20 min. The aqueous layer was loaded on iodixanol gradients in a Type 70.1 rotor and centrifuged for 2 h at 52000 rpm. The rAAV sample was collected and buffer exchanged with 1x PBS supplemented with 5% Sorbitol and 0.1M NaCl (0.25M NaCl final) using an Amicon 100kda filter.

For all surgeries, mice received pre-operative analgesia with meloxicam (10 mg/kg, subcutaneous) and buprenorphine (0.1 mg/kg, subcutaneous). Anaesthesia was induced with 5% isoflurane and maintained at 2–2.5% throughout the procedure. Mice were positioned in a stereotactic frame, and bupivacaine (1.25 mg/ml) was applied as a local anaesthetic to the scalp. Eye ointment was applied to prevent corneal drying, and body temperature was maintained at 37 °C using a feedback-controlled heating pad.

For experiments involving 3-month-old mice, a combined craniotomy and AAV injection surgery was performed at 2 months of age. A custom titanium headplate was fixed to the skull using dental cement (Tetric EvoFlow), and a 4 mm-diameter craniotomy was made over the somatosensory cortex using a high-speed drill. AAVs were injected directly into the exposed cortex using pulled glass capillaries and a Nanoject injector at a rate of 0.1 μl/min (400 nl per construct). The stereotactic coordinates relative to bregma were: ML = ±1.7 mm, AP = −1.4 mm, and DV = −1.1 mm (depth from skull surface). The capillary was left in place for 10 min after injection to allow for diffusion. Cranial windows were constructed by bonding a 4 mm round coverglass to a 5 mm outer coverglass using optical adhesive (Norland). The assembled window was implanted over the craniotomy and sealed using veterinary adhesive (Gluture) and dental cement. Post-operative care included oral meloxicam (5 mg/kg in custard) every 24 hours for 3 days.

For experiments involving 8-month-old mice, AAVs were injected at 2 months of age via a small burr hole drilled above the somatosensory cortex using the same stereotactic coordinates and injection parameters as described above. The scalp was closed with surgical glue, and mice were returned to their home cage. At 7 months of age, mice underwent a second surgery to implant a cranial window for *in vivo* two-photon imaging. The craniotomy and window implantation followed the same protocol as described for the 3-month-old cohort, including pre-and post-operative care.

Surgeries were performed 3–4 weeks prior to experimental readouts. Mice were either 3 or 8 months old at the time of imaging or tissue collection, depending on the study.

### Two-photon imaging

Mice were anaesthetised with 1.2% isoflurane, and eye ointment was applied to prevent corneal drying. Body temperature was maintained at 37 °C using a heating pad throughout imaging. Two-photon imaging was performed using a resonant/galvo/galvo microscope equipped with two GaAsP detectors and a Coherent Chameleon Discovery TPC dual-laser system. Imaging was controlled using ScanImage software (Vidrio Technologies) running in MATLAB (MathWorks, Natick, MA, USA). Images were acquired at 512 × 512 pixel resolution and 5 Hz using a Nikon 25× water-immersion objective (NA 1.1). Imaging sessions lasted 10 minutes to record spontaneous calcium activity in astrocytes at depths of 150–300 μm below the pial surface. GCaMP and mCherry signals were acquired at excitation wavelengths of 920 nm and 1100 nm, respectively.

Two-photon calcium imaging data were analysed using ImageJ/Fiji (https://imagej.nih.gov) and custom-written scripts in MATLAB following a previously published protocol². Fluorescence time series were first motion-corrected, and regions of interest (ROIs) were manually segmented based on signal from both GCaMP and mCherry channels. For astrocytes, ROIs encompassed the soma and proximal processes. ΔF/F₀ traces were computed by normalising the fluorescence signal (F) to a baseline (F₀), defined as the mean of the lowest 10% of fluorescence values within each trace. All traces were linearly detrended prior to event detection. Calcium transients were defined as ΔF/F₀ signals that exceeded 2x the standard deviation of the baseline noise. We then quantified the number of transients and ΔF/F₀ amplitude for each group.

### Immunohistochemistry

Mice were euthanised via intraperitoneal injection of pentobarbital (60 mg/kg), followed by transcardial perfusion with ice-cold phosphate-buffered saline (PBS) and subsequently with 4% paraformaldehyde (PFA). Brains were carefully extracted and post-fixed in 4% PFA for 24 h at 4 °C, after which 40 μm-thick coronal sections were prepared using a vibratome. Sections were stored at −20 °C in cryoprotectant solution (30% ethylene glycol, 30% glycerol, 40% PBS) until further use.

Free-floating brain sections were blocked for 1h at room temperature in blocking buffer (PBS containing 0.1% Triton X-100 and 5% goat serum). Immunofluorescence was performed using the following primary antibodies, each diluted 1:500 in blocking buffer: rabbit anti-LAMP1 (Abcam, ID ab208943), rat anti-LAMP2 (Abcam, ID ab13524), rabbit anti-Cathepsin D (Abcam, ID ab75852), guinea pig anti-S100β (Synaptic Systems, ID 287004), rabbit anti-TRPML1 (Proteintech Europe, ID 15291-1-AP), chicken anti-GFAP (Aves Labs, ID SKU GFAP), and mouse anti-Aβ (clone 82E1; Tecan, ID JP10323). Sections were incubated with primary antibodies overnight at 4 °C, followed by three washes in PBS. Secondary antibodies, also diluted 1:500 in blocking buffer, included Alexa Fluor 594 goat anti-rat IgG (H+L) (Invitrogen, ID A-11007), Alexa Fluor 488 goat anti-guinea pig IgG (H+L) (Invitrogen, ID A-11073), Alexa Fluor 647 goat anti-chicken (Invitrogen, ID A32933) and Alexa Fluor 555 goat anti-rabbit IgG (H+L) (Invitrogen, ID A-21428). Sections were incubated with secondary antibodies overnight at 4 °C, washed in PBS, and mounted using Dako fluorescence mounting medium (Agilent Technologies, USA).

Images were acquired using a Leica TCS SP8 confocal inverted microscope equipped with LAS X software. Image analysis was performed using ImageJ/FIJI software. Quantification of lysosomal expression was based on the area of overlap (double-positive signal) between each lysosomal marker (LAMP1, LAMP2, Cathepsin D, and TRPML1) and S100β. Quantification of FIRE-pHLy probe consisted of calculating the ratio of mTFP/mCherry signals. For amyloid pathology, the area stained by the Aβ-specific antibody 82E1 was quantified, along with the size of individual plaques, measured in square micrometres.

### Statistical analyses

Details on the number of animals and samples analysed are provided in the results section and figure legends. Statistical analyses were performed using GraphPad Prism (GraphPad Software, San Diego, CA, USA), MATLAB (MathWorks, Natick, MA, USA), and R (R Foundation for Statistical Computing, Vienna, Austria). Data were first assessed for normality using the Shapiro–Wilk test. Where appropriate, data were log-transformed to meet the assumptions of parametric testing. Depending on the distribution and experimental design, either nested two-sample t-tests or nested one-way ANOVAs with Holm-Sidak correction for multiple comparisons were used to compare two or more groups. For nested data that did not meet parametric assumptions, we applied the aligned rank transform (ART) method, a non-parametric approach suitable for factorial and mixed-effects models. For comparisons between two groups, ART ANOVA was performed with group as a fixed effect and subject as a random effect. For analyses involving more than two groups, ART ANOVA was followed by planned pairwise post hoc comparisons with Holm-Sidak correction for multiple testing.

## Author Contributions

D.S., B.D.S., and I.L.A.C. conceived and designed the study. D.S., P.D., C.C., M.J.T., M.D., and V.F.T.M. performed the experiments. D.S. and V.F.T.M. were responsible for animal colony establishment and post-surgical care. D.S. carried out data acquisition and analysis. M.S. provided key experimental materials and contributed technical guidance on study design. D.S., B.D.S., and I.L.A.C. wrote the manuscript. All authors contributed to reviewing and editing the manuscript and approved the final version. B.D.S. and I.L.A.C. supervised the work and acquired funding.

## Data Availability

The data supporting the findings of this study are available from the corresponding author upon reasonable request.

## Ethics approval declaration

All experimental procedures were conducted under the authority of UK Home Office project licence PP4652483 in accordance with the Animals (Scientific Procedures) Act 1986, and were approved by the Francis Crick Institute Animal Welfare and Ethical Review Body (AWERB).

