## Supplemental figures for "TRPML1 loss drives lysosomal calcium failure and astrocyte dysfunction across Alzheimer’s Disease progression"

[3.] VIB Center for brain and disease research and Leuven Brain Institute, KU Leuven, Leuven, Belgium

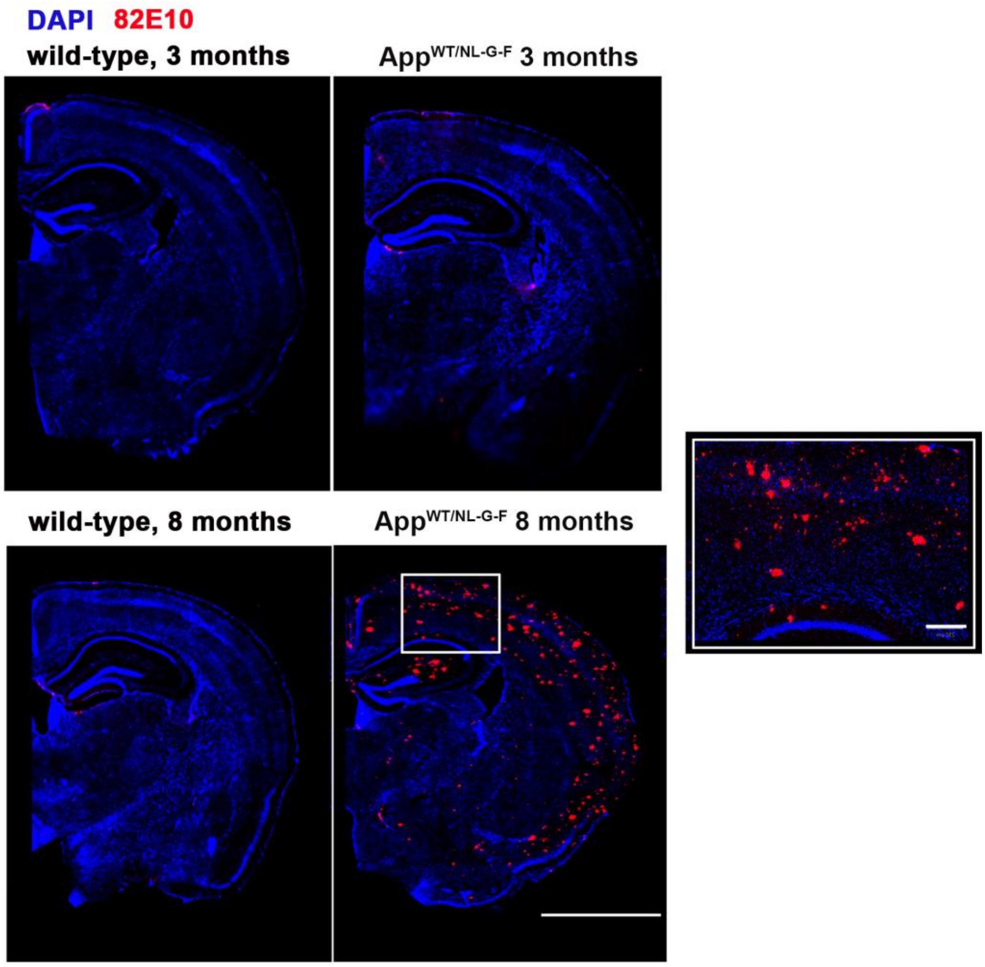


**<u>Figure S1:</u> Time course of amyloid plaque deposition in heterozygous App$^{WT/NL-G-F}$ mice.**

Representative immunohistochemistry images showing 82E1 staining for amyloid-β in cortex and hippocampus of App$^{WT/NL-G-F}$ mice at 3 and 8 months of age. No amyloid-β signal was detected at 3 months, confirming the absence of plaque pathology at this pre-plaque stage, whereas robust amyloid deposition was evident by 8 months. Inset: higher-magnification view of cortical region from an 8-month-old App$^{WT/NL-G-F}$ mouse illustrating dense plaque accumulation.

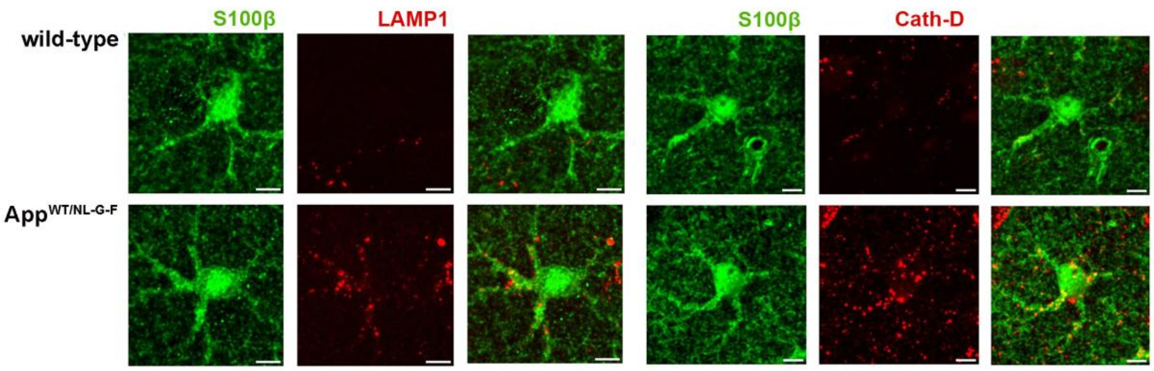


**<u>Figure S2:</u> Early changes in lysosomal markers in AD astrocytes.** Immunohistochemistry images showing LAMP1 and cathepsin D expression in astrocytes (S100β) from wild-type (N = 5 mice, 8 cells per mouse) and $App^{WT/NL-G-F}$ mice (N = 6 mice, 8 cells per mouse).

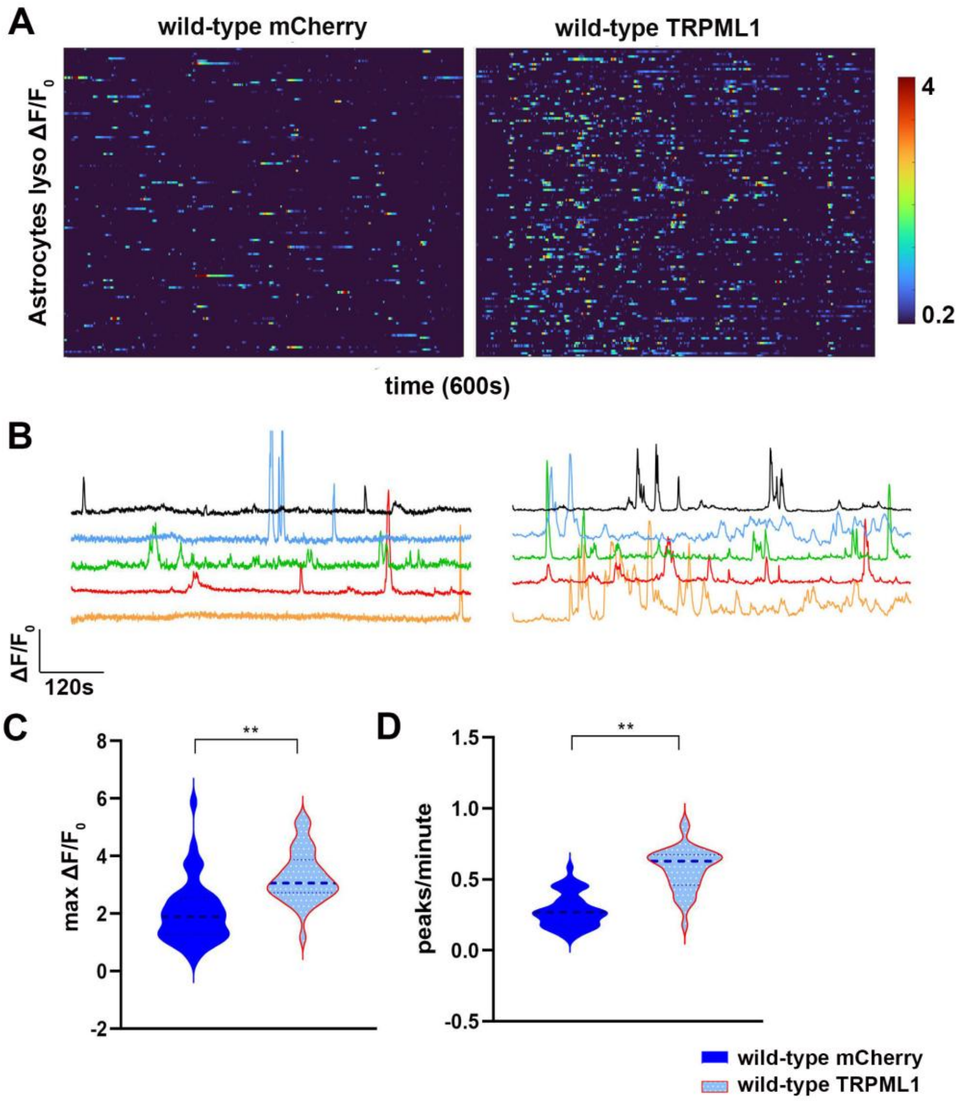


<u>Figure S3:</u> **TRPML1 expression increases lysosomal calcium release in healthy astrocytes.**

**A)** *In vivo* two-photon calcium imaging of wild-type (WT) mice expressing AAV-GFAP-LAMP1-GCaMP6f co-injected with either AAV-GFAP-mCherry or AAV-GFAP-TRPML1. Heatmaps show lysosomal calcium transients ($\Delta F/F_0$) over 600 s for individual astrocytes. **B)** Five representative calcium traces are displayed per group. **C-D)** Quantification of $\Delta F/F_0$ amplitude and calcium event frequency (mean ± SEM) is shown for mCherry- and TRPML1-expressing groups (*N* = 4 mice/group, 25–39 astrocytes per mouse). Amplitude: 1.85 ± 0.15 (mCherry) vs. 3.28 ± 0.15 (TRPML1), $p = 0.0081$. Frequency: 0.27 ± 0.018 (mCherry) vs. 0.61 ± 0.023 (TRPML1), $p = 0.0095$. Statistical test: ART ANOVA.

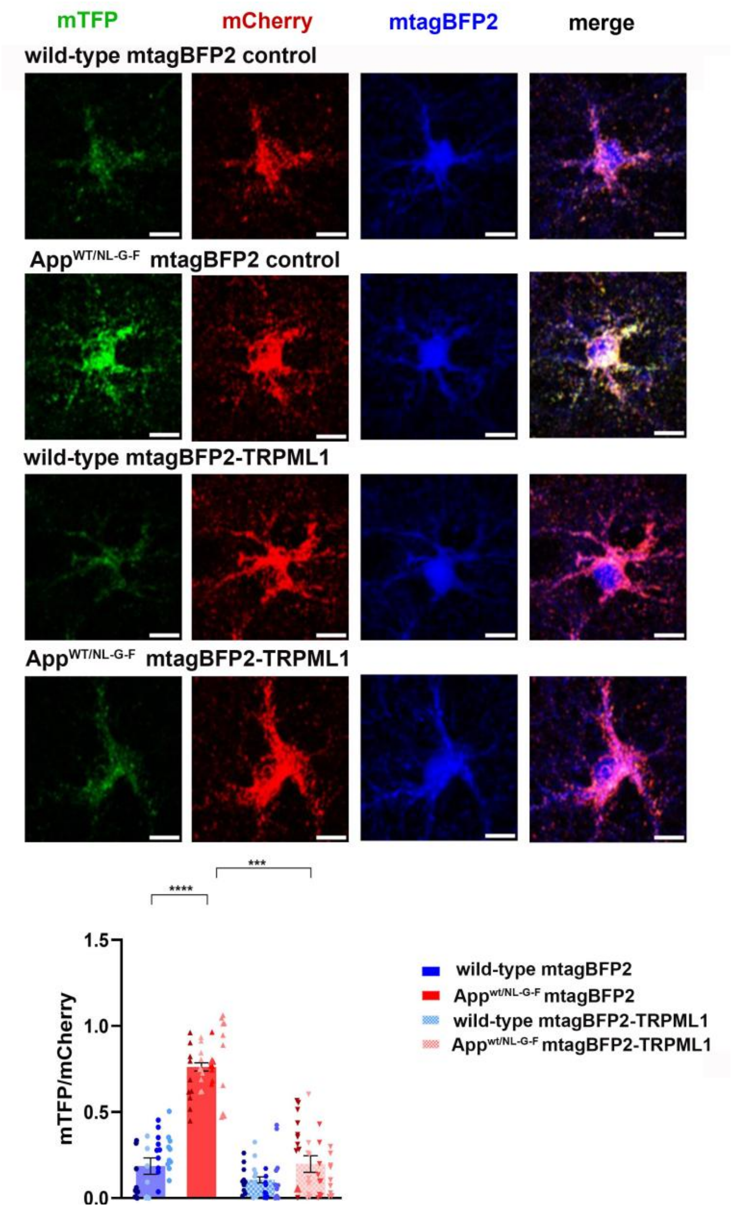


**<u>Figure S4:</u> Early TRPML1 expression normalises astrocytic lysosomal acidity in App$^{WT/NL-G-F}$ mice.** Representative images showing expression of the AAV-GFAP-FIRE-pHLy probe in astrocytes expressing AAV-GFAP-mtagBFP2 (control) or AAV-GFAP-mtagBFP2-TRPML1 (N = 4 mice /experimental group, 9-11 cells per mouse). Quantification of the mTFP/mCherry ratio (mean ± SEM): WT mtagBFP2, 0.18 ± 0.04; App$^{WT/NL-G-F}$ mtagBFP2, 0.76 ± 0.04; WT mtagBFP2-TRPML1, 0.11 ± 0.03; App$^{WT/NL-G-F}$ mtagBFP2-TRPML1, 0.19 ± 0.05. Significant differences: WT vs App$^{WT/NL-G-F}$ mtagBFP2, *p* =

0.0007; App$^{WT/NL-G-F}$ mtagBFP2 vs. mtag-TRPML1, *p*=0.0007. Statistical test: ART ANOVA with Holm-Sidak correction.

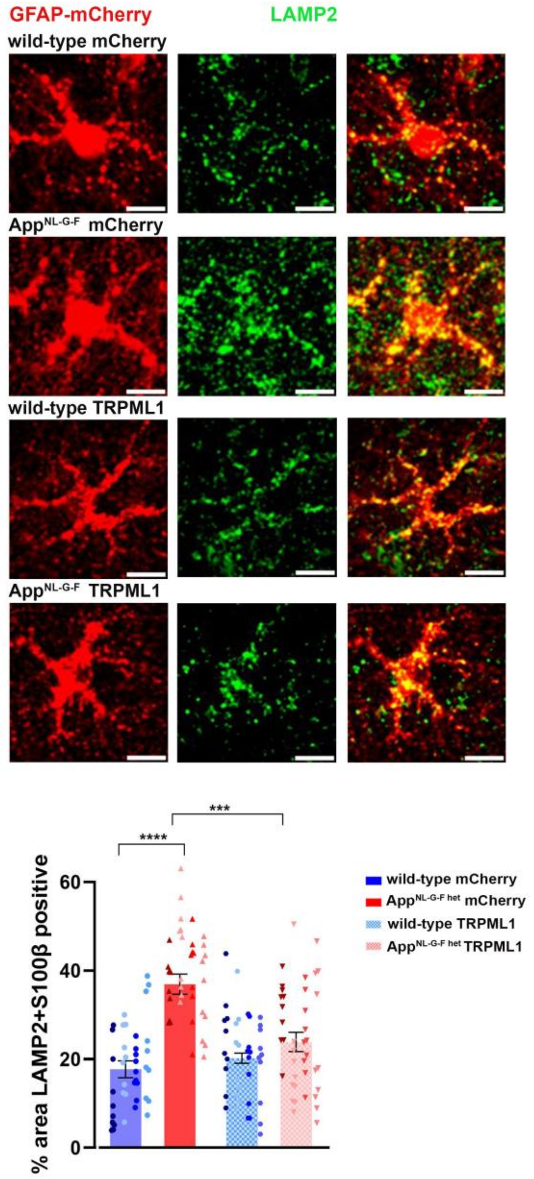


<u>Figure S5:</u> **Early TRPML1 expression in astrocytes normalises lysosomal LAMP2 in App$^{WT/NL-G-F}$ mice.** Immunohistochemistry images showing LAMP2 expression in astrocytes expressing either AAV-GFAP-mCherry or AAV-GFAP-TRPML1-mCherry (N = 4 mice/experimental group, 10 cells per mouse). Quantification is shown as % area of pixels double-positive for LAMP2 and mCherry (mean ± SEM). Values: WT mCherry, 18 ± 2.4; App$^{WT/NL-G-F}$ mCherry, 37 ± 2.6; WT TRPML1, 19 ± 2.6; App$^{WT/NL-G-F}$ TRPML1, 24 ± 3.0.

Significant differences: WT vs. $App^{WT/NL-G-F}$ mCherry, $p < 0.0001$; $App^{WT/NL-G-F}$ mCherry vs. TRPML1, $p<0.0001$. Statistical test: Nested one-way ANOVA with Holm-Sidak correction.
